# TMEM174 Deficiency Reduces Longevity by Promoting Phosphate-Driven Vascular Calcification

**DOI:** 10.64898/2026.04.09.716713

**Authors:** Jose G Miranda, Judith Blaine, Makoto Miyazaki

## Abstract

**Background:** Dysregulation of phosphate homeostasis contributes to reduced longevity and vascular complications in chronic kidney disease and aging. This study investigates the role of TMEM174, a proximal tubule-specific protein, in regulating the phosphate co-transporter NPT2A and its subsequent impact on lifespan and vascular health.

**Methods:** TMEM174 knockout (KO) mice (C57BL6/J and DBA/2J) were fed diets with varying phosphate concentrations (0.6% vs. 1.2%). In OKP cells, TIRF and FRET microscopy, alongside immunoprecipitation, were used to identify the TMEM174 protein regions essential for NPT2A binding and endocytosis.

**Results:** TMEM174 KO mice exhibited significantly shorter lifespans than wild-type controls. High phosphate diets exacerbated vascular calcification, stiffness, and mortality, while low phosphate diets rescued these phenotypes. In vitro, TMEM174 siRNA blocked PTH-induced NPT2A endocytosis, increasing its apical membrane retention. FRET and biochemical assays revealed that the C-terminal region of TMEM174 is essential for its association with NPT2A. While intact TMEM174 and N-terminal mutants (TMEM174ΔN) facilitated NPT2A degradation, C-terminal deletions (TMEM174ΔC) failed to associate with or degrade NPT2A.

**Conclusions:** TMEM174 is a critical regulator of phosphate homeostasis and longevity. The C-terminal region of TMEM174 is specifically required for NPT2A endocytosis and degradation, identifying it as a potential therapeutic target for managing phosphate-related vascular complications.

## Introduction

The regulation of phosphate balance is critical to maintaining systemic homeostasis, and disruptions in these processes are tightly linked to the progression of chronic kidney disease (CKD), ageing, metabolic disorders, and cardiovascular complications. Among the transporters mediating renal phosphate handling, sodium-dependent phosphate transporter type IIa (NPT2A)—encoded by the *SLC34A1* gene—plays a central role in phosphate reabsorption in the proximal tubule of the kidney^1-3^. NPT2A is regulated by a complex interplay of hormonal and intracellular signals, including parathyroid hormone (PTH) and fibroblast growth factor 23 (FGF23), which modulate its membrane localization, degradation and activity ^4-7^. Mutations or dysregulation of NPT2A have been implicated in conditions such as hyperphosphatemia, hypophosphatemia, nephrolithiasis, and CKD progression, making it a critical node for understanding renal phosphate transport and systemic mineral balance ^8-11^.

Gene-targeted disruptions in phosphate homeostasis such as the Klotho-FGF23 axis in mice cause aging-like symptoms including a very short life span, while klotho overexpression extends the longevity of mice. In addition, phosphate lowering interventions such as low phosphate diet feeding and NPT2A inhibition rescue the aging-like phenotypes of klotho and FGF23 knockout (KO) mice^11-14^. More interestingly, levels of serum phosphate inversely correlate with the average life span of mammals. High serum phosphate is associated with all causes of mortality in humans^12,15,16^. These observations suggest that phosphate is a critical factor for determining the life span of mammals.

Emerging interest has focused on Transmembrane Proteins (TMEM), a relatively understudied membrane protein whose expression patterns and putative signaling roles suggest involvement in cellular stress responses and immune-related pathways. While the precise physiological functions of many TMEM proteins remain to be fully elucidated, transcriptomic and proteomic analyses have indicated the differential regulations in disease states, including renal and metabolic disorders^17,18^. Recently, we and another research group have reported that Transmembrane Protein 174 (TMEM174) is involved in phosphate homeostasis by regulating NPT2A ^19,20^. TMEM174 is a type III transmembrane protein that is specifically expressed in the brush border membrane of renal proximal tubular epithelial cells^19^. TMEM174 knockout mice developed severe hyperphosphatemia and vascular calcification due to increasing levels of NPT2A protein in the kidney ^19^. In addition, TMEM174 inhibition blocked FGF23- and PTH-mediated degradation of NPT2A *in vivo* and in cultured proximal tubular cells. However, the precise mechanism by which TMEM174 regulates the degradation of the NPT2A protein is not fully understood.

In this study, to advance understanding of the physiological and biological role of TMEM174, we examine whether 1) TMEM174 deficiency shortens the life span of mice, 2) varying dietary phosphate concentrations affect the life span and vascular calcification in TMEM174 KO mice and 3) which part of the TMEM174 protein is critical in regulating the endocytosis of NPT2A.

## Results

### TMEM174KO mice exhibit shorter life spans and vascular dysfunction, dependent on phosphate levels

We previously reported that TMEM174 is a novel regulator in phosphate homeostasis ^19^. Since circulating phosphate is considered a major factor in determining the life span of mammals, we tested whether TMEM174 deficiency affects longevity of mice on two mouse genetic backgrounds, C57BL/6J and DBA/2J. As shown in Figure 1, TMEM174 deficiency drastically shortens the life span of mice on both backgrounds. On the DBA/2J background, the average life span of TMEM174 KO mice was 76 days and 185 days for male and female mice, (Figure 1A), respectively, whereas the average life span of C57BL/6J TMEM174KO male mice was 256 days,, which is much longer than DBA/2J TMEM174KO mice (Figure 1B). We next examined whether the shorter life span of TMEM174KO mice is associated with vascular dysfunctions such as vascular stiffness and calcification. At 8 weeks of age, mice were subjected to aortic pulse wave velocity analysis using an Indus Doppler Flow Velocity System. On both genetic backgrounds, the aortic pulse wave velocity (aPWV) was greater in TMEM174 KO mice compared to wild-type counterparts. Consistent with the shorter life span, the aPWV of DBA TMEM174KO mice was significantly higher than C57BL6 TMEM174KO and wild-type mice (Figure 1C). Histological analysis with von Kossa staining and aortic calcium analysis showed that TMEM174 KO mice on both C57BL6 and DBA/2J backgrounds developed aortic calcification (Figure 1D-1F). Importantly, vascular calcification in DBA TMEM174KO mice was drastically more severe than in C57BL6 TMEM174KO mice (Figure 1D-1F). We analyzed biomedical parameters that are related to kidney function and phosphate homeostasis. On both C57BL6 and DBA/2J backgrounds, TMEM174 deficiency did not alter glomerular functions as creatine levels, and eGFR was not different. TMEM174KO mice, however, had significantly higher levels of serum phosphate on both C57BL6 and DBA/2J backgrounds (Table 1). More importantly, DBA2/J TMEM174KO mice developed more severe hyperphosphatemia than C57BL6/J TMEM174KO mice (Table 1). In addition, TMEM174KO mice on both genetic backgrounds had significantly higher levels of the phosphate regulating hormones FGF23 and PTH than wild-type counterparts (Table 1). Interestingly, FGF23 levels in C57Bl6/J TMEM174KO mice were higher than those in DBA2J TMEM174KO mice. Levels of serum calcium, cholesterol, triglycerides, and KIM-1 are comparable between wild-type and TMEM174KO mice on both genetic backgrounds (Table 1). Regardless of TMEM174 genotype, serum cholesterol levels were higher and KIM-1 levels were lower on the DBA/2J genetic background (Table 1).

**Table 1.** Serum parameters of C57Bl/6J and DBA/2JTMEM174 KO mice fed chow diet.

|  | C57BL/6J-WT | C57BL6-TMEM174KO | DBA2/J-WT | DBA2/J-TMEM174KO | WT vs KO | DBA vs C57 |
| --- | --- | --- | --- | --- | --- | --- |
| Creatinine (μg/ml) | 1.02 ± 0.02 | 1.06 ± 0.02 | 0.81 ± 0.04 | 0.88 ± 0.03 | NS | NS |
| GFR (μl/min) | 272.12 ± 16.71 | 265.5 ± 27.63 | 249 ± 20.08 | 241.875 ± 17.2 | NS | NS |
| Phosphorus (mg/ml) | 4.80 ± 0.32 | 8.44 ± 0.23<sup>a</sup> | 4.64 ± 0.31 | 14.39 ± 0.18<sup>b</sup> | p<0.0001 | NS |
| FGF23 (pg/ml) | 518.42 ± 2821.21 | 18769.50 ± 67.81<sup>a</sup> | 460.98 ± 6.82 | 8058.75 ± 799.51<sup>b</sup> | p<0.0001 | NS |
| PTH (pg/ml) | 62.07 ± 14.49 | 1318.97 ± 163.21<sup>a</sup> | 421.62 ± 48.56 | 2079.45 ± 312.69<sup>b</sup> | p<0.0001 | p<0.0001 |
| Calcium (mg/ml) | 10.23 ± 0.64 | 9.13 ± 0.19 | 9.22 ± 0.25 | 9.20 ± 0.17 | NS | NS |
| Cholesterol (mg/ml) | 93.75 ± 14.15 | 95.91 ± 6.82 | 381.64 ± 28.39 | 222.12 ± 57.65 | NS | p<0.0001 |
| Triglyceride (mg/ml) | 161.96 ± 9.04 | 122.10 ± 14.73 | 118.48 ± 10.53 | 121.05 ± 11.37 | NS | NS |
| KIM-1 (pg/ml) | 63.92 ± 14.49 | 72.6371 ± 13.6803 | 37.77 ± 4.75 | 41.28 ± 3.62 | NS | p<0.0001 |

Serum was collected from eight-week old male (N=8) and female (N=3) mice fed a chow diet,.
NS: Not significant; <sup>a</sup>P<0.05 vs C57BL/6J-WT; <sup>b</sup>P<0.05 vs DBA2/J-WT
Two-way ANOVA was used for statistical analysis.

**Figure 1.**
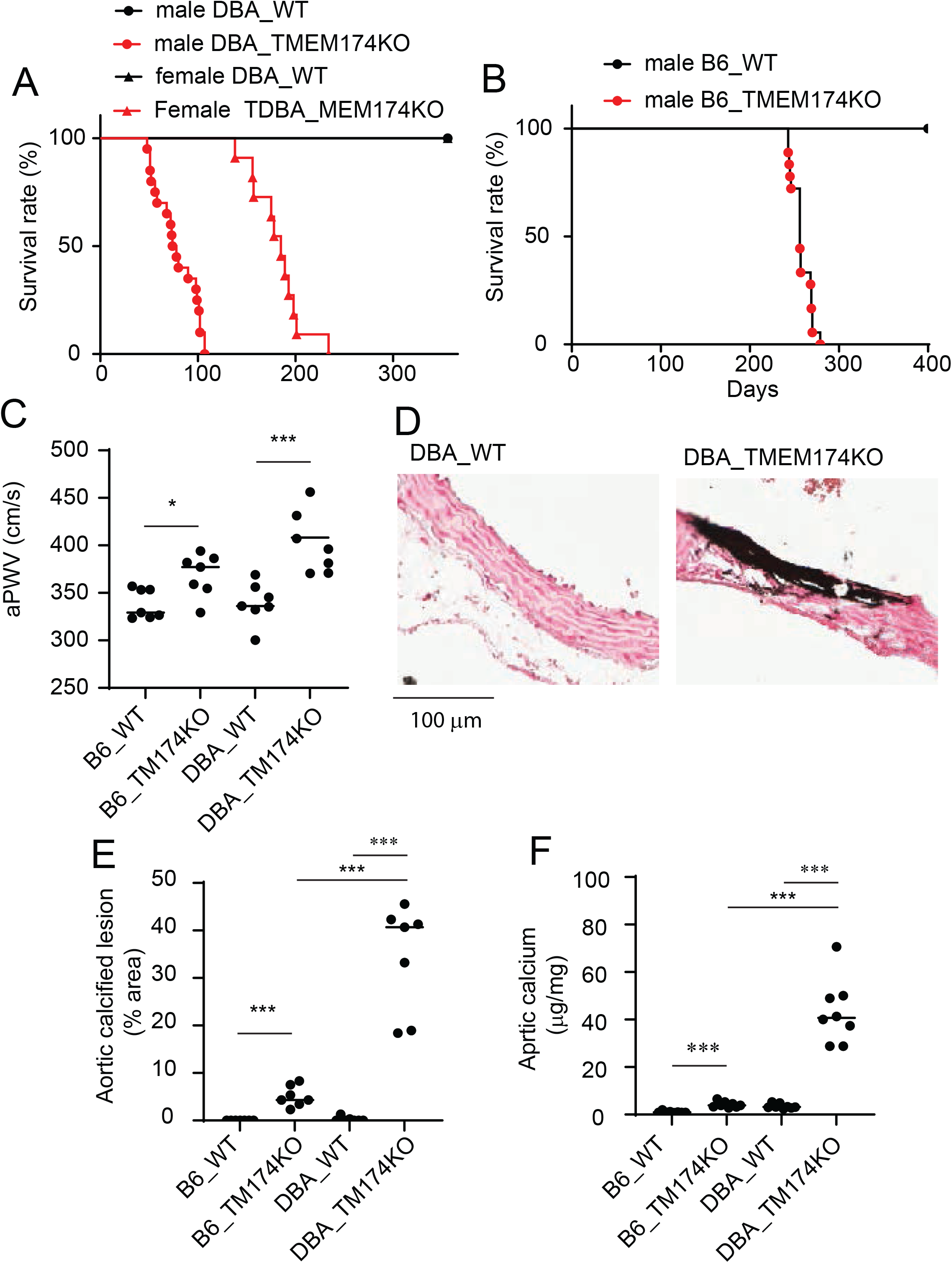
TMEM174 KO mice on both C57BL6/J and DBA2/J backgrounds had short life spans associated with vascular stiffness and calcification. A) Survival rate of C57BL6/J TMEM174KO male mice (N=24) and B) survival rate of DBA2/J TMEM174 KO male and female mice (N=24). C) Aortic pulse wave velocity (aPWV) of TMEM174KO male mice. aPWV was analyzed at 8 weeks of age using an Indus pulse wave velocity system. D) Representative pictures of aortic calcification. Vascular calcification was analyzed with von Kossa staining using aortic arches. Aortas were isolated from 8-week-old male mice. E) % area of calcified lesions in aortic arches. F) Aortic calcium content in C57BL6/J and DBA2/J TMEM174KO mice.

To examine whether regulating phosphate homeostasis is critical for longevity in TMEM174KO mice, at 8 weeks of age animals were fed different concentrations of dietary phosphate: 1.2% phosphate (TD.85349) for a high phosphate diet, 0.6% phosphate (TD.84122) for a normal phosphate diet, and 0.1% phosphate (TD.85010) for a low phosphate diet. A DBA/2J background was used as this is more sensitive to TMEM174 deficiency. The high phosphate diet extremely shortened the longevity of TMEM174KO mice. The average life span was 12 days on the high phosphate diet, whereas TMEM174KO mice on the normal phosphate diet survived for an average of 61 days. The low phosphate diet on the other hand significantly extended the survival rate of TMEM174KO mice. Dietary phosphate concentrations did not affect the longevity of wild-type mice. To examine if dietary phosphate affects vascular functions, we analyzed aPWV and vascular calcification after 1 week of high, normal, and low phosphate feeding, and after 4 weeks of normal and low phosphate feedings, as all the TMEM174KO mice on the high phosphate diet died within 3 weeks (Figure 2A). TMEM174KO mice on high and normal phosphate diets had greater aPWV than wild-type counterparts, whereas aPWV of TMEM174 and wild-type mice were comparable on the low phosphate diet (Figure 2B). High phosphate and normal phosphate diets induced vascular calcification in TMEM174KO mice (Figure 2C-2E). The high phosphate diet induced more severe vascular calcification than the normal phosphate diet (Figure 2C-2D). More importantly, the low phosphate diet blocked the development of vascular calcification in TMEM174KO mice. None of the diets induced vascular calcification in wild-type mice(Figure 2C-2D). Consistently, in TMEM174KO mice, the high phosphate diet significantly increased aortic calcium content compared to the normal phosphate diet (Figure 2E). We next examined whether dietary phosphate affects the biomedical parameters related to phosphate homeostasis and kidney function. High phosphate diet exacerbated hyperphosphatemia in TMEM174KO mice whereas low phosphate diet attenuated hyperphosphatemia in TMEM174KO mice, compared to normal phosphate diet (Figure 3A). Levels of FGF23 are highest in TMEM174KO mice fed the high phosphate diet followed by TMEM174KO mice fed the normal phosphate diet. Low phosphate diet on the other hand normalized the extremely higher levels of FGF23 in TMEM174KO mice (Figure 3B). PTH levels were significantly higher in TMEM174KO mice fed high phosphate and normal phosphate diets compared to wild-type counterparts, whereas low phosphate diet normalized levels of PTH (Figure 3C). Low phosphate diet also reduced levels of serum phosphate, FGF-23 and PTH in wild-type mice compared to normal phosphate diet. Regardless of diets and genotypes, levels of serum creatinine, calcium and KIM-1 were comparable (Figure 3D-F).

**Figure 2.**
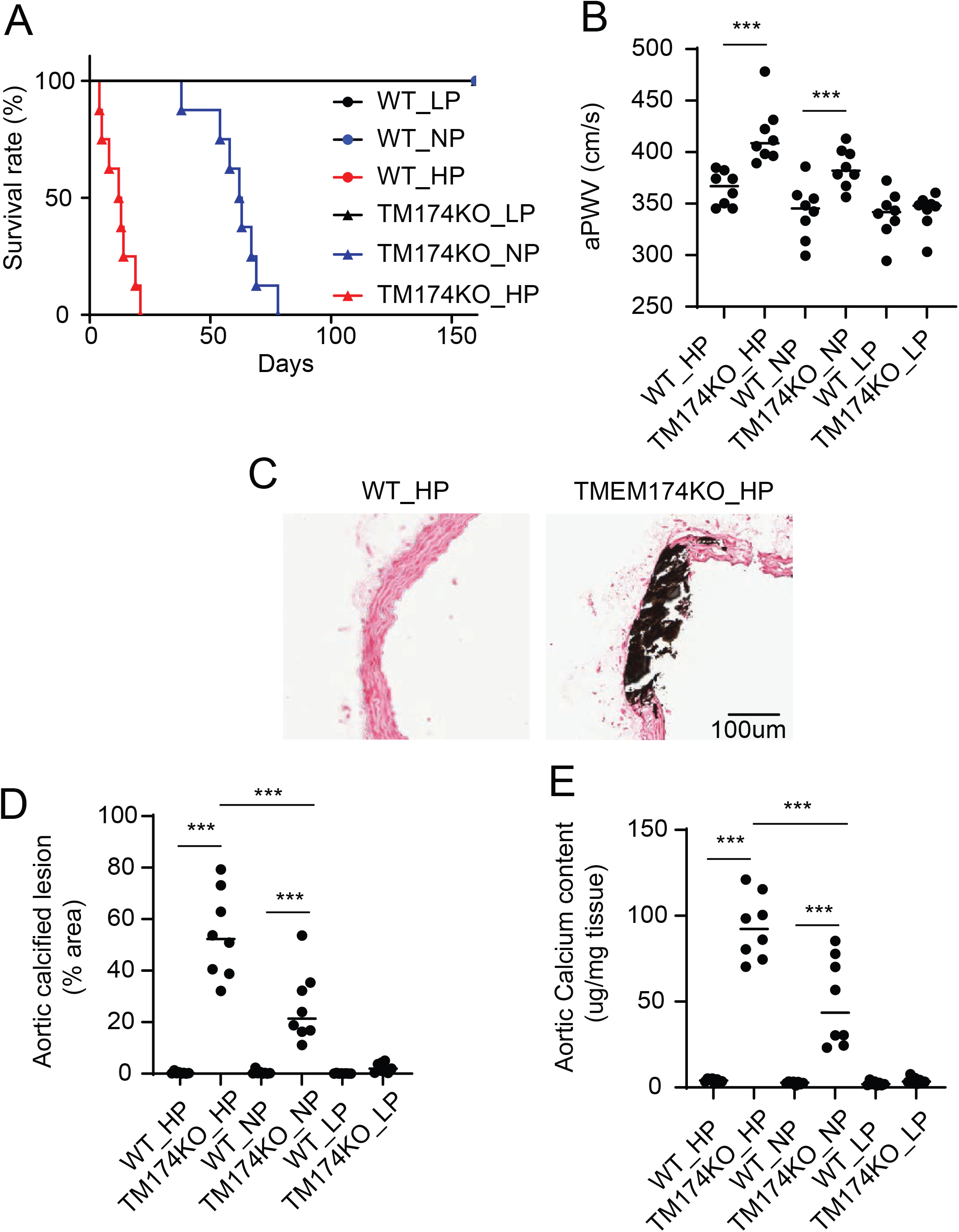
High phosphate diet shortens the life span of DBA2/J TMEM174KO mice, whereas low phosphate diet extends the life span. A) Survival rate of male DBA2/J TMEM174KO mice (6-week-old) fed various concentrations (HP: 1.2%, NP: 0.6% and LP: 0.1%) of dietary phosphate. B) Aortic pulse wave velocity (aPWV) of TMEM174KO mice fed various concentrations of dietary phosphate. aPWV was analyzed 7 days after the start of the feeding using an Indus pulse wave velocity system. C) Representative picture of an aortic arch with von Kossa staining. Feeding with high phosphate diet induces vascular calcification in TMEM174KO mice, whereas feeding with the low phosphate diet attenuates vascular calcification. Aortas were isolated after 10 days of the feeding. D) % area of calcified lesions in aortic arches, and E) aortic calcium content in C57BL6/J and DBA2/J TMEM174KO mice

**Figure 3.**
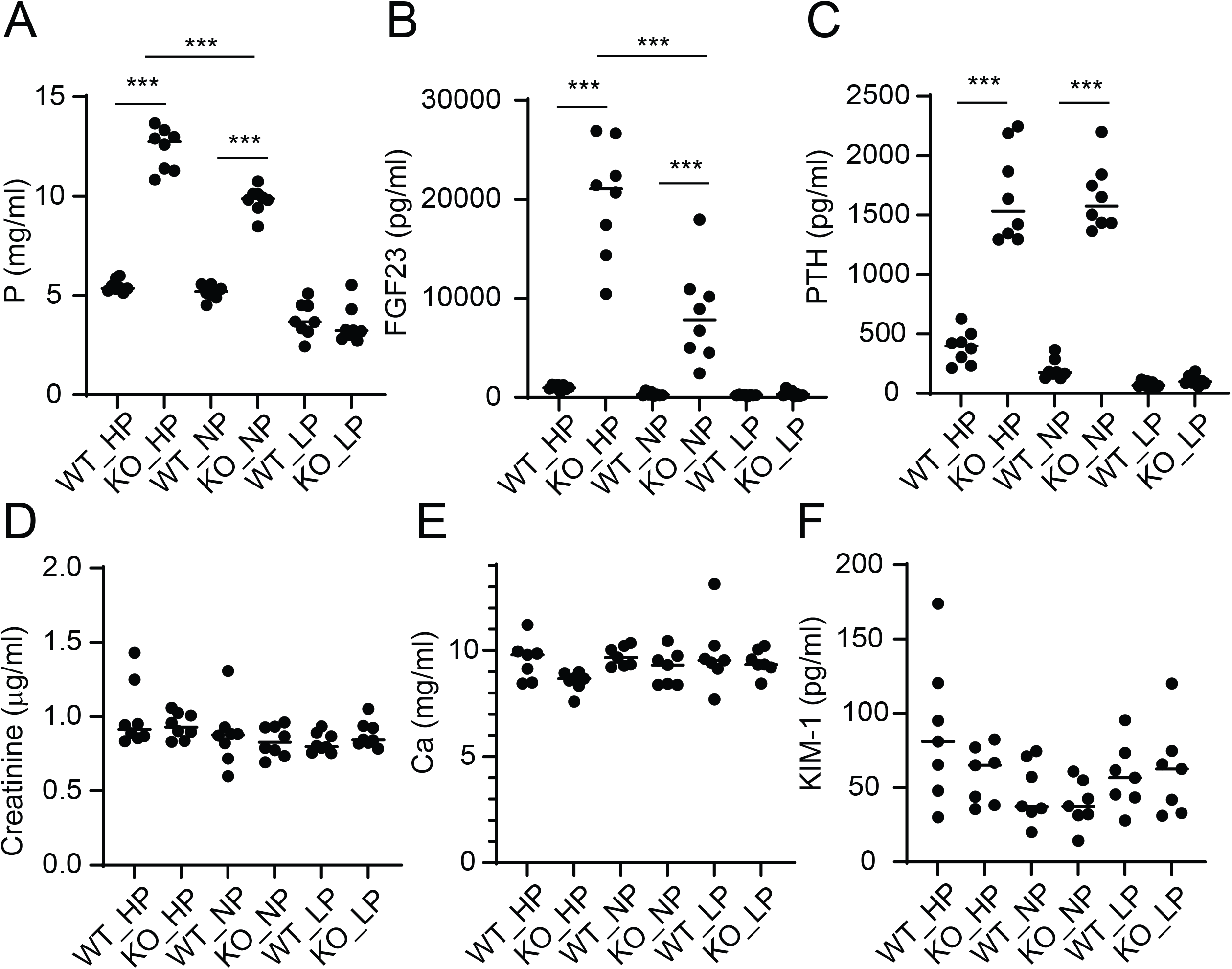
Biomedical parameters related with phosphate homeostasis and kidney function. Levels of serum A) phosphorus, B) FGF-23, C) PTH, D) creatinine, E) calcium and F) KIM-1 in male DBA2/J TMEM174KO mice fed various concentrations of dietary phosphate. Mice were euthanized 7 days after the start of the feeding.

We next examined whether dietary phosphate affects NPT2A expression in the proximal tubules using immunoblot and immunofluorescence analysis. Similar to C57BL6J background, under high phosphate, DBA2/J TMEM174KO mice had extremely higher levels of NPT2A in the renal BBM. TMEM174KO mice fed a low phosphate diet had higher NPT2A protein in the renal BBM than wild-type mice fed a low phosphate diet (S. Figure 1).

### Knock-down of TMEM174 in OKP cells blocks NPT2A endocytosis via the parathyroid hormone

We have previously shown that TMEM174 deficiency significantly increased levels of NPT2A in both renal brush boarder membranes *in vivo* and in proximal tubule cells *in vitro* ^19^. In addition, TMEM174 deficiency completely blocked PTH- and FGF23-mediated reduction of NPT2A in mice and renal proximal tubule cells. However, the mechanism by which TMEM174 destabilizes NPT2A has not been fully elucidated. To obtain an understanding of the mechanism, we have performed a series of experiments to examine which part of the TMEM174 protein is critical in regulating NPT2A endocytosis. We first examined whether TMEM174 deficiency retains NPT2A protein in the plasma membrane in proximal tubule cells. Like our previous report using total protein lysates^19^, PTH treatment significantly reduced NPT2A protein in the membrane fraction of OKP cells treated with control siRNA, as expected (Figure 4A&4B). In addition to TMEM174 siRNA treatment drastically increasing NPT2A in the membrane under basal conditions, PTH-mediated NPT2A endocytosis was blocked by TMEM174 knockdown (Figure 4A&4B). Levels of membrane TMEM174 protein were not changed upon PTH treatment (Figure 4A). To modulate and track the fluorescence of NPT2A and TMEM174, we used different constructs and transiently transfected them into OKP cells. For NPT2A, one construct used the yellow fluorescent protein (YFP) and the other construct used mScarlet, which is a red shifted fluorescent protein. YFP and mScarlet were placed at the N-terminal end of NPT2A. For TMEM174, moxCerulean-3 (mC3) and green fluorescent protein (GFP) were placed at the C-terminal of TMEM174. Two separate fluorescent proteins were used because the total internal reflection fluorescent (TIRF) microscope at our disposal only contains a green laser and a red laser. For the YPF-tagged NPT2A we used wide-field microscopy. Seven days after splitting and seeding OKP cells into 35mm glass cover imaging dishes, cells were transfected with siRNA primers specifically used for knocking-down TMEM174. These siRNA primer transfections were done a minimum of three separate times to ensure TMEM174 knock-down. 48 hours prior to imaging, cells were transfected with NPT2A tagged with either YFP or mScarlet. Cells were imaged using a Nikon TiE widefield microscope or a Zeiss Elyra P.1 PALM/STORM super resolution microscope. Consistent with the immunoblot analysis of endogenous NPT2A protein, Figures 4C and 4D demonstrate that PTH induced the degradation of EYFP-NPT2A in the presence of TMEM174, whereas PTH-induced NPT2A degradation occurred minimally in the absence of TMEM174. As shown in Figures 4E and 4F, when performing the same experiment under the same conditions using inverted TIRF microscopy to look at the apical membrane more precisely as previously described^4^, we observed similar results compared to wide-field microscopy. When TMEM174 is knocked-down there is less decay of m-Scarlet-NPT2A in the apical membrane compared to when TMEM174 is present. This demonstrates that TMEM174 is responsible for the regulation of NPT2A endocytosis. TMEM174 protein in the apical membrane was not affected by PTH treatment (Figure 4G-4J). Both wide-field and TIRF microscopy analysis show that there is no decay over time in mC3 fluorescence and GFP over the course of PTH treatment. (Figure 4) This confirms that NPT2A tagged fluorescence decays over time as a function of PTH.

**Figure 4.**
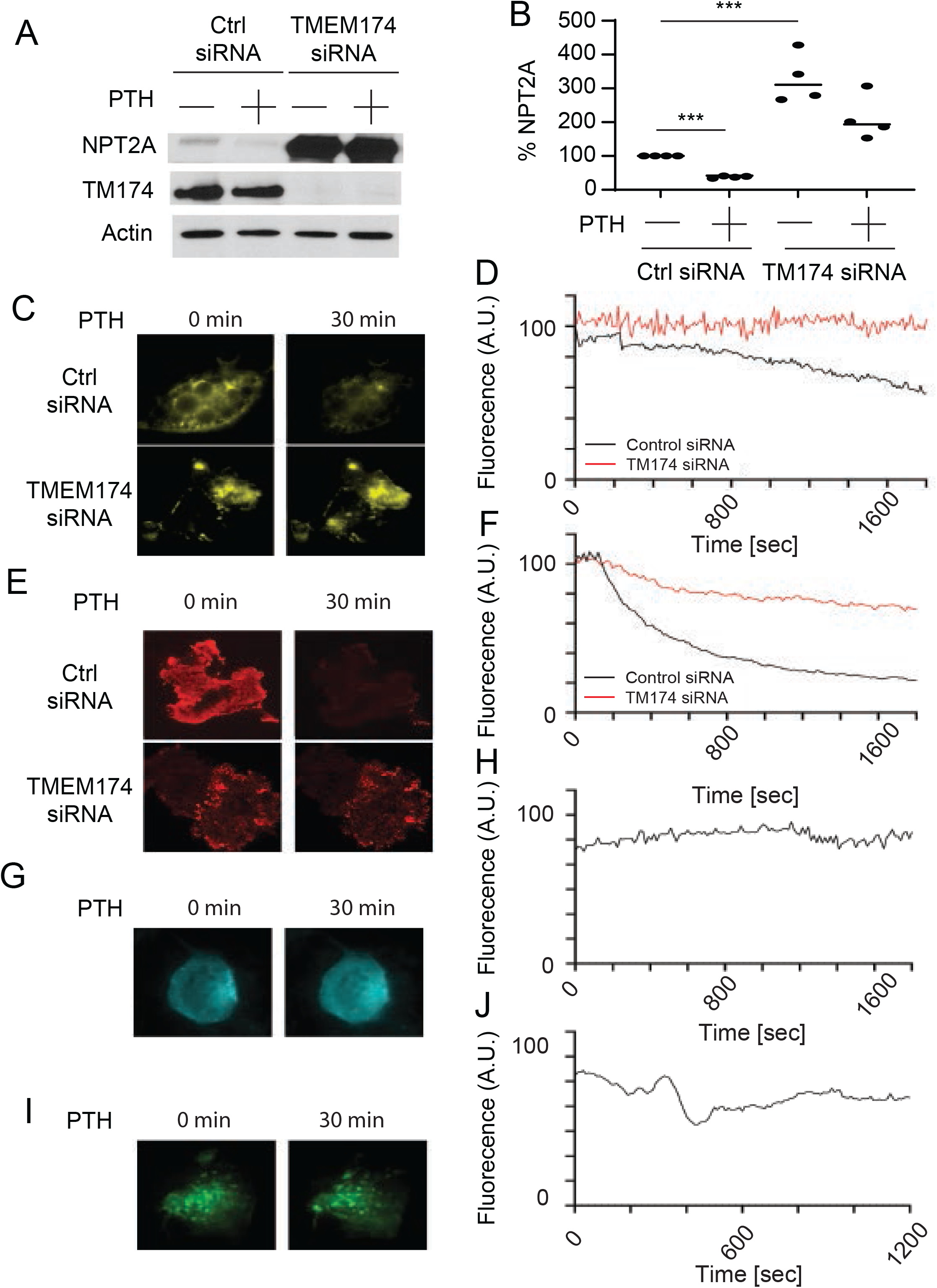
TMEM174 is involved in PTH-mediated NPT2A endocytosis. A) Immunoblot analysis and B) densitometry analysis of the NPT2A immunoblot analysis in OKP cells treated with TMEM174 siRNA and PTH. OKP cells were treated with 20nM TMEM174 siRNA or scramble control siRNA in the presence of RNAi Max Reagent. After 2 days at 100% confluency, OKP cells were treated with 10nM PTH for 1 hour. Plasma membrane fraction was isolated with sequential ultracentrifuge. C) Representative image of widefield microscopy of OKP cells treated with tagged YFP (EYFP-NPT2A) and PTH. D) The graph represents the widefield fluorescent microscopy of EYFP-NPT2A in the presence of TMEM174 siRNA and PTH. E) Representative image of inverted TIRF microscopy of OKP cells expressing NPT2A tagged with mScarlet (mScarlet-NPT2A) treated with PTH and TMEM174 siRNA. F) The graph represents the inverted TIRF microscopy of OKP cells expressing mScarlet-NPT2A treated with PTH and TMEM174 siRNA. G) The representative images of the widefield microscopy showing the fluorescence over time of TMEM174 tagged with moxCerulean3 (TMEM174-mC3) in OKP cells treated with PTH. H) The graph represents the widefield microscopy of TMEM174-mC3 fluorescent proteins in OKP cells treated with PTH. I) The representative images of the TIRF microscopy show the fluorescence over time of TMEM174 tagged with EGFP (TMEM174-EGFP) in OKP cells treated with PTH. J) The graph represents the TIRF microscopy of TMEM174-EGFP fluorescent proteins. Cells were treated with TMEM174 siRNA for three rounds prior to imaging. Twenty-four hours before imaging, cells were placed in FBS deficient media. Widefield and TIRF imaging was done over the course of 30 minutes, acquiring images every 10 seconds and 15 seconds for widefield and inverted TIRF, respectively.

### The C-terminal end of TMEM174 is responsible for NPT2A regulation

We next tested which region of the TMEM174 protein is responsible for the regulation of NPT2A. We removed the N-terminal and C-terminal ends that span the channel outside of the plasma membrane. Figure 5A shows the schematic of TMEM174 and specifically which region was removed. To test the interaction between TMEM174 and NPT2A, immunoblot blot analysis was performed after pulling down using FLAG-tagged TMEM174 full-length (wt), TMEM174ΔN (N-terminal truncation), and TMEM174ΔC (C-terminal truncation). Figure 5C shows that samples transfected with TMEM174wt and TMEM174ΔN were able to pull down endogenous NPT2A protein with a FLAG monoclonal antibody, while the samples with TMEM174ΔC expression did not pull down the NPT2A protein. The co-immunoblot of NPT2A shows that it is pulled down with both flag-TMEM174wt and flag-TMEM174ΔN but not flag-TMEM174ΔC, suggesting that NPT2A binds to the C-terminal end of TMEM174. To further analyze whether the C-terminal end of TMEM174 is crucial in the regulation of NPT2A, we performed microscopic FRET analysis. Figure 5D shows OKP cell images of the FRET channel that have been transfected with YFP-NPT2A with TMEM174wt-mC3, TMEM174ΔN-mC3, or TMEM174ΔC-mC3. The warmer the color, the higher the FRET ratio, and the cooler the color, the less FRET is occurring. Additionally, because the C-terminal region of TMEM174 is regulated by NPT2A, the top and middle panels show that the FRET ratio decreases in a time dependent manner after PTH is added. Most importantly, the lower panel with TMEM174ΔC shows the FRET ratio is low and there is no change over time after PTH is added. This shows that NPT2A is not binding with TMEM174 and corroborates the data shown in the Western blot. Figure 5E shows the experiments performed on the initial FRET ratio and demonstrates that the initial FRET ratio is the same in both TMEM174wt and TMEM174ΔN, and they are not statistically significant. However, there is a significant FRET ratio difference comparing TMEM174ΔC to both TMEM174wt and TMEM174ΔN. Figure 5F shows that the FRET ratio decreases in a time-dependent manner as a function of PTH for both TMEM174wt and TMEM174ΔN co-transfected with NPT2A, but there is no change in the FRET ratio when using TMEM174ΔC with NPT2A. Plotting the end point of these experiments shows that the FRET ratio decreases between NPT2A and either TMEM174wt or TMEM174ΔN compared to that of TMEM174-ΔC, fig 5G. This data further corroborates that NPT2A binds to the C-terminal end of TMEM174 and PTH induces the disassociation between NPT2A and TMEM174. To determine whether 1) TMEM174 mutants affect PTH-induced NPT2A endocytosis and 2) PTH alters TMEM174 proteins, as well as to confirm photobleaching does not affect the fluorescent proteins, we plotted fluorescence in a time-dependent manner as a function of PTH. Figures 5H and 5J show that there is minimal to no decay of mC3 attached to all three variants of TMEM174. PTH reduced EYFP-NPT2A signal in the presence of TMEM174wt and TMEM174ΔN, whereas in the presence of TMEM174ΔC, EYFP-NPT2A signals show no decay due to PTH treatment (Figure 5K and 5L). This data demonstrates and solidifies that the C-terminal end of TMEM174 is responsible for the regulation of NPT2A. To confirm the results from the microscopic analysis, we performed immunoblot analysis to determine whether the C-terminal end of TMEM174 is crucial in the regulation of NPT2A. TMEM174 knockdown increased levels of NPT2A in OKP cells in the presence and absence of PTH. More importantly, the restoration of TMEM174 by human TMEM174wt and TMEM174ΔN recovered TMEM174 knockdown-mediated NPT2A induction in the absence and presence of PTH, whereas human TMEM174ΔC retrieval did not affect TMEM174 knockdown (S. Figure 2A). To further test this interaction and how it affects NPT2A binding to TMEM174, we made several C-terminal truncations, removing the last 43 or 86 amino acids of TMEM174. We tested these short variants of TMEM174 against full-length TMEM174, and one in which the entire C-terminal region was deleted. Supplemental Figure 2 shows images of all channels, CFP, YFP, and FRET under basal condition. The average resting FRET ratio of these pairs shows that full-length TMEM174 has a higher resting FRET ratio compared to the TMEM174ΔC truncated mutants, suggesting that the first 43 amino acids of the C-terminal region of TMEM174 are very important in bringing NPT2A in close proximity to TMEM174.

**Figure 5.**
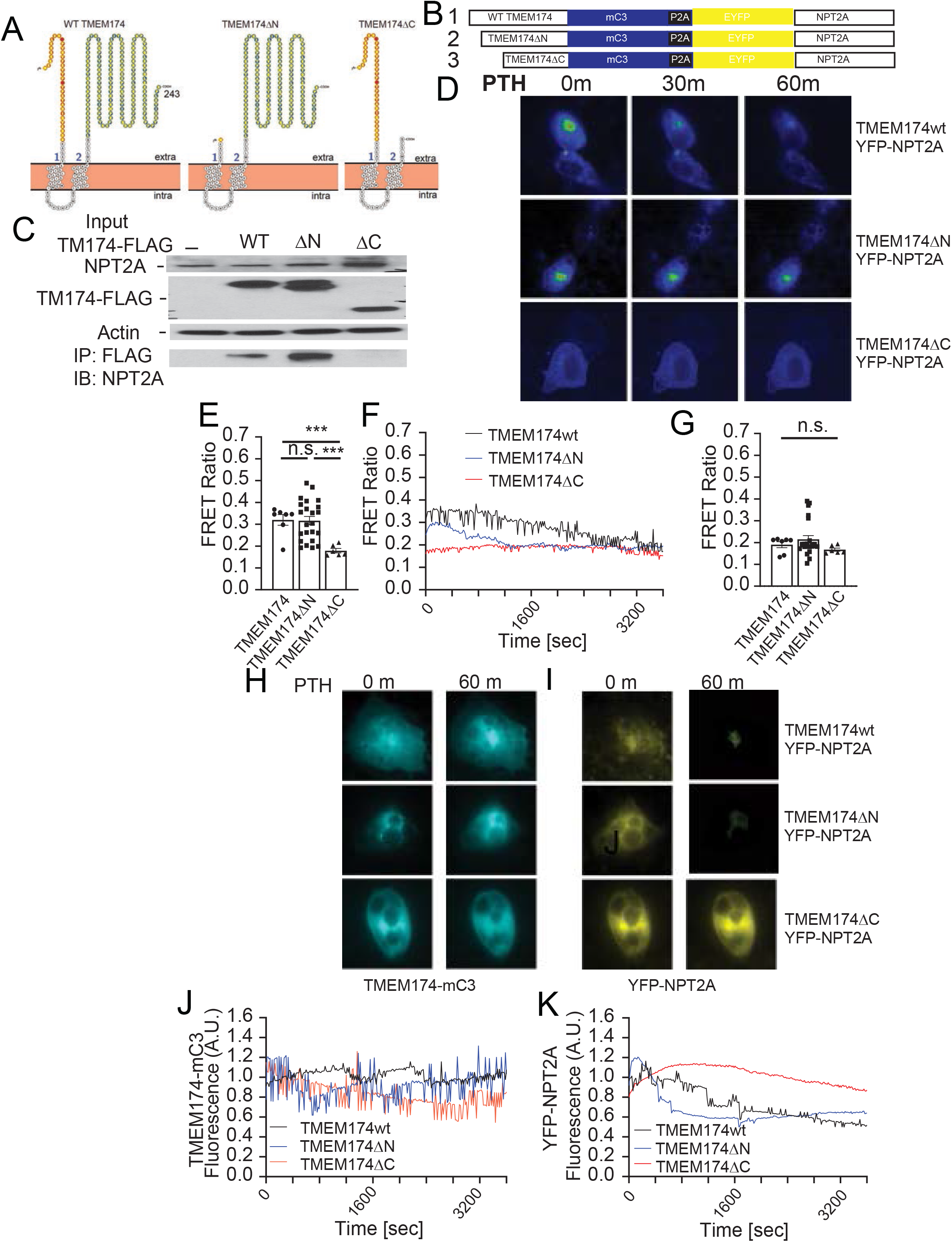
The TMEM174 C-terminal end is critical in the regulation of NPT2A endocytosis. A) The schematic of TMEM174 and truncations performed that were used with NPT2A and the schematic of which terminal region the FLAG tag or fluorescent protein was placed on. The NPT2A N-terminal end was tagged with EYFP, whereas the TMEM174 C-terminal end was tagged with FLAG or mC3. B) Co-immunoprecipitation of endogenous NPT2A. TMEM174 wild-type-FLAG (TMEM174wt), TMEM174-FLAG lacking the N-terminal end (TMEM174ΔN) or TMEM174-FLAG lacking the C-terminal end (TMEM174ΔC) were transiently overexpressed in OKP cells. Total cell protein lysate was immunoprecipitated with mouse anti-FLAG monoclonal antibody. The immunoprecipitated proteins were immunoblotted with rabbit anit-NPT2A antibody. C) The dual expression vectors for FRET analysis. To simultaneously express EYFP-NPT2A and TMEM174-mC3 mutants, a ribosomal skipping sequence, P2A was inserted between NPT2A and TMEM174. D) The representative image showing the FRET ratio of TMEM174 and NPT2A combinations during PTH treatment. E) The initial FRET ratio between EYFP-NPT2A and TMEM174-mC3 mutants. F) Time course of FRET analysis during PTH treatment. This demonstrates that the C-terminal end is involved in the regulation of NPT2A as a function of PTH. G) The end FRET ratio between EYFP-NPT2A and TMEM174-mC3 mutants during PTH treatment using widefield microscopy. J) The mC3 fluorescence signal of TMEM174 and K) the YFP fluorescence signal of NPT2A during PTH treatment in the presence of TMEM174 mutants using widefield microscopy.

## Discussion

In the present study, we demonstrate that TMEM174 is a critical determinant of systemic phosphate homeostasis, longevity, and vascular integrity, and we identify the C-terminal region of TMEM174 as essential for PTH-induced NPT2A endocytosis. Using two genetic backgrounds and dietary phosphate manipulation, we show that TMEM174 deficiency leads to severe hyperphosphatemia, accelerated vascular calcification, and a markedly shortened life span in a phosphate-dependent manner. Mechanistically, live-cell imaging, FRET, and co-immunoprecipitation analyses reveal that the C-terminal domain of TMEM174 mediates interactions with NPT2A and is required for hormone-stimulated internalization and degradation. Together, these findings establish TMEM174 as a previously unrecognized regulator linking renal phosphate transport to organismal aging and vascular pathology.

A major physiological insight from this study is that loss of TMEM174 dramatically shortens mouse life spans, and that this phenotype is strongly modulated by dietary phosphate. High-phosphate intake rapidly exacerbated mortality and vascular stiffness in TMEM174-deficient mice, whereas phosphate restriction normalized biochemical abnormalities, prevented vascular calcification, and substantially extended survival. These data provide direct *in vivo* evidence that the reduced longevity caused by TMEM174 deficiency is largely mediated through disordered phosphate metabolism rather than primary renal failure, as glomerular function remained preserved. The findings align with prior genetic models in which disruption of phosphate-regulating pathways accelerates aging-like phenotypes and lowering phosphate rescues survival ^13,14,21^. Our results extend this concept by identifying TMEM174 as a proximal tubular factor required for adaptive phosphate excretion and lifespan maintenance under physiological conditions.

We also observed strain-dependent differences in disease severity, with DBA/2J TMEM174-deficient mice showing more severe hyperphosphatemia, vascular calcification, and shorter life spans than C57BL/6J counterparts. These data suggest that genetic modifiers such as gene mutations in Abcc6 that regulate systemic levels of inorganic pyrophosphate and in CYP24A1 that catabolize vitamin D on the DBA/2J and C57BL/6J genomes^22-24^, respectively, influence susceptibility to phosphate toxicity when TMEM174 function is impaired. Interestingly, despite higher FGF23 levels in C57BL/6J TMEM174-deficient mice, phosphate levels and vascular pathology were worse in DBA/2J animals, implying differential target-organ responsiveness to phosphaturic hormones between strains. Such variation may help explain heterogeneity in phosphate-related vascular disease across populations and warrants future investigation into downstream signaling or renal transporter regulation.

At the cellular level, our imaging studies clarify the mechanism by which TMEM174 controls NPT2A abundance. TMEM174 knockdown prevented PTH-induced removal of NPT2A from the apical membrane, leading to persistent transporter retention. FRET and co-immunoprecipitation experiments demonstrated that full-length TMEM174 and the N-terminal truncation mutant bind NPT2A, whereas C-terminal deletion abolishes this interaction. Moreover, only constructs containing the C-terminal region restored PTH-mediated NPT2A degradation in TMEM174-deficient cells. These complementary approaches indicate that the extracellular C-terminal domain of TMEM174 is necessary for physical association with NPT2A and for hormone-triggered endocytosis. The observation that progressive C-terminal truncations reduced basal FRET efficiency further suggests that the proximal portion of the C-terminus contains key residues required for transporter docking.

Our data support a model in which TMEM174 acts as an accessory membrane component that enables hormonal signaling to access the NPT2A endocytic machinery. Under physiological conditions, PTH promotes dissociation of NPT2A from TMEM174 followed by internalization and degradation, thereby reducing phosphate reabsorption. In the absence of TMEM174 or its C-terminal interaction domain, NPT2A remains stabilized at the brush border membrane and becomes resistant to PTH-induced trafficking, resulting in sustained phosphate uptake and systemic overload. This mechanism is consistent with the *in vivo* phenotype of TMEM174-deficient mice, which exhibit elevated renal NPT2A, hyperphosphatemia, and vascular calcification despite increased circulating FGF23 and PTH.

The link between TMEM174, phosphate burden, and vascular disease observed here has important translational implications. Hyperphosphatemia is a major driver of vascular calcification and mortality in CKD and aging populations ^25-27^. Our findings suggest that impaired TMEM174 function could represent a previously unrecognized contributor to phosphate retention and vascular pathology. Conversely, strategies that enhance TMEM174 activity or mimic its C-terminal interaction with NPT2A may represent novel approaches to promote phosphaturia and limit vascular calcification. The strong rescue by dietary phosphate restriction in TMEM174-deficient mice also underscores the therapeutic importance of phosphate control in conditions of impaired renal phosphate handling.

Several limitations should be considered. First, although we demonstrate phosphate-dependent life span changes, we cannot exclude additional TMEM174 functions that may influence aging or vascular biology independently of NPT2A. Second, the precise molecular partners linking the TMEM174 C-terminus to the endocytic machinery remain unidentified as TMEM174 does not directly bind to the NPT2A scaffold protein, NHERF1^19,20^. Third, our mechanistic studies were performed in proximal tubule cells and require confirmation *in vivo* at endogenous expression levels. Finally, whether TMEM174 variation contributes to human phosphate disorders or vascular calcification remains unknown.

In summary, this study identifies TMEM174 as a critical regulator of renal phosphate transport, vascular calcification, and life spans, and establishes the C-terminal domain of TMEM174 as essential for PTH-induced NPT2A endocytosis. These findings reveal a new molecular axis controlling phosphate homeostasis and provide mechanistic insight into how disrupted proximal tubular phosphate handling can drive systemic aging-related pathology. Further characterization of TMEM174 signaling and its interaction partners may uncover new therapeutic targets for hyperphosphatemia-associated diseases.

## Materials and Methods

### Mice

All mouse experiments were performed in compliance with the University of Colorado Anschutz Medical Campus International Animal Care and Use Committee (IACUC). Mice were provided with water and food *ad libitum* and were housed in an environment with adequate temperature control with a 12h light/dark cycle. C57BL/6J and DBA/2J mice were obtained from the Jackson Laboratory. TMEM174KO mice on C57BL6/J and DBA2/J backgrounds were generated as previously described^19^. Mice were fed a chow diet (2020, Inotiv, Madison) or diets with different concentrations of phosphate: low (0.1%) phosphate diet (TD.85010, Inotiv, Madison, WI), normal (0.6%) phosphate diet (TD.84122, Inotiv), and high (1.2%) phosphate diet (TD.85349, Inotiv). The diets were otherwise matched on calcium, magnesium, sodium, protein, fat, and vitamin D content. Feeding of the semi-purified diet was started at 6 weeks of age.

#### Biomedical parameter analysis

Serum calcium (Ca) and phosphorus (P) were analyzed using commercial kits with colorimetric assays (Pointe Scientific). Serum FGF23 and PTH were analyzed with ELISA kits from Quidel (60-6300 and 60-2350, respectively) whereas KIM-1 was analyzed with an ELISA kit from R&D systems (MKM100). Serum creatinine was analyzed using LC-MS/MS as previously described^28-32^.

#### Cell Culture

Opossum proximal kidney tubule (OKP) cells were grown in Dulbecco’s Modified Eagle Medium/F-12 (D-MEM/F12, Gibco). Medium was supplemented with 10% (v/v) fetal bovine serum (FBS, Fisher Scientific), 100 U/mL penicillin, and 100 µg/mL streptomycin (Corning). Cells were incubated at 37 °C in 5% CO_2_ in a humidified controlled environment, and the medium was changed every 2-3 days. OKP cells were seeded onto 35 mm glass bottom imaging dishes (No 1.5 cover glass). Seven days post seeding, after cells were already differentiated, 2 µg of pcDNA3.1 containing YFP-NPT2A, GFP-NPT2A, Scarletl-NPT2A, TMEM174-mCerulean3/YFP-NPT2A, TMEM174ΔN-mC3/YFP-NPT2A, TMEM174ΔC-mC3/YFP-NPT2A, TMEM174ΔC86-mC3/YFP-NPT2A, or TMEM174ΔC43-mC3/YFP-NPT2A were transiently transfected using 3 µL of Lipofectamine 2000 (Invitrogen), gently mixed with 120 µL of OPTI-Mem and let sit for 15 minutes in the tissue culture hood. This was followed by the addition of 1.5 µg of DNA and mixed gently, followed by a 45 min incubation in the tissue culture hood. Prior to adding the transfection cocktail to OKP cells, the media was removed from dish, the cells were briefly washed with PBS pH 7.4, and 2.5 mL of fresh DMEM/F12 media was added before returning the dishes to the incubator.

### Live-cell microscopy and TIRF and FRET imaging and analysis

Forty-eight to 72 hours post-transfection, cells were imaged using a modified imaging solution (130 mM NaCl, 5 mM KCl, 1.2 mM MgCl_2_ · Heptahydrate, 2.5 mM CaCl_2_ · Dihydrate, 11 mM D(+)-glucose, and 30 mM HEPES). An Eclipse-TiE wide-field microscope (Nikon) with a Nikon Plan Fluor 40x/1.30 oil immersion objective was used for OKP cells. An X-Cite Series 120PC-Q filter switcher and shutter controller were used (Excelitas Technologies). Images were acquired at 1 frame/15 seconds for 30 – 60 minutes using a BFLY-U3-23S6M-C monochrome camera, with µManager Software to operate the system. For confocal microscopy a Zeiss LSM780 microscope with a C-Apochrome 40x/1.20 W Korr FCS M27 objective using the 405 nm, 488 nm, and 561 nm wavelength lasers was used on fixed samples. For total internal reflection fluorescence (TIRF), a Zeiss Elyra P.1 (PALM/STORM) microscope with a α-Plan apochromat 100x/NA 1.46 Oil DIC M27 Elyra objective using 488 nm and 561 nm wavelength lasers at 1% power was used. Images were acquired at 1 frame/15 seconds for 15-30 minutes to minimize bleaching using an Andor iXon+ 897 EMCCD camera. For mCerulean3 and YFP experiments, the FRET ratio (^FRET^/_mCerulean3_ = FRET ratio) was calculated by dividing the FRET signal (YFP channel signal upon CFP excitation) by mCerulean3 (CFP channel signal upon CFP excitation) and plotted over time. For YFP, GFP, or mScarlet signal, regions of interest were normalized to 1.0. For analysis we used Student’s unpaired t-test.

### Glomerular filtration rate (GFR) analysis

The dialyzed FITC-inulin solution was sterile filtered and injected into the retro-orbital plexus (2 μl/g body weight) during brief isoflurane anesthesia. At 3, 7, 10, 15, 20, 40 and 60 minutes after the injection, blood was collected from a small tail nick the end of the tail. After centrifugation the fluorescence was determined using a fluorospectrometer.

#### BBM vesicle isolation

Renal brush border membrane vesicles (BBM) were prepared from mouse kidneys by the Ca^2+^ precipitation method, as described previously^4,5,19^. OKP cells were grown to confluence in 100mm dishes. 24 hours before the experiment, the cells were placed in DMEM medium containing 0.1% BSA to synchronize them. Cells were incubated with 10 nM PTH for 1 hour. After treatment, the cells were washed in ice cold PBS and scraped into isolation buffer (15 mmol/l Tris (pH7.4), 300 mmol/l mannitol, 5 mmol/l ethylene glycol tetra acetic acid, and one Mini-Complete tablet (Roche) on ice. The cells were then homogenized by aspirating 30 times through a 23-gauge needle. CaCl_2_ was added to a final concentration of 100mM, and the homogenate was shaken on ice for 20 minutes. The homogenate was centrifuged at 2500 x g at 4 C for 15 minutes. The supernatant was removed and spun at 41,000 x g for 30 minutes. The final pellet was resuspended in RIPA buffer.

#### Immunoblot analysis

Brush border membrane vesicles (BBMVs) were prepared from mouse kidney and OKP cells by the Ca^2+^ precipitation method, as described previously^4,5^. BBMVs protein samples were subjected to SDS/PAGE. The separated proteins were transferred by electrophoresis to PVDF membranes (Immobilon-P; Millipore). The membranes were treated with affinity-purified anti-mouse or opossum NPT2A antibody (Biomatik), anti-opossum or mouse TMEM174 antibody (Biomatik), anti-FLAG M2 (Sigma-Aldrich F1804) and anti-Actin (1:2000, Proteintech 66009). Goat anti-rabbit or anti-mouse IgG-horseradish peroxidase conjugate was utilized as the secondary antibody, and signals were detected using Supersignal ECL reagents (Thermo Fisher). The uncropped pictures are shown in the supplement.

#### Co-immunoprecipitation

OKP cells were transfected with pcDNA3.1 containing TMEM174-FLAG wild-type and TMEM174-mutants using Lipofectamine 2000. Cell lysates were prepared using co-immunoprecipitation buffer (150 mM NaCl, 1% Nonidet P-40, 50 mM Tris, pH 8.0). Five hundred μg of lysate was incubated with 40 μl of anti-DYKDDDDK G1 Affinity Resin (GenScript) overnight at 4 °C. Resins were washed 5 times with co-immunoprecipitation buffer. Immunoprecipitated proteins were analyzed using immunoblot analysis with anti-opossum NPT2A and Veriblot for IP detection reagent (Abcam, ab131366)^19,33^.

#### Immunohistochemical analysis

Immunostaining analysis of mouse kidney sections (Zyagen) was performed as described previously^19^. For immunofluorescence, the section was incubated with affinity-purified anti-TMEM174 monoclonal antibody (Biomatik, this antibody was generated with a full-length mouse TMEM174 protein as the antigen), anti-mouse NPT2A polyclonal antibody and Lotus Tetragonolobus Lectin (LTL, a renal apical membrane marker, Vector Laboratories FL-1321-2) overnight at 4°C. Immunoreactivity was detected using Alexa Fluor 488-conjugated anti-mouse IgG (Thermo Fisher Scientific A-11001) or Alexa Fluor 568-conjugated anti-rabbit IgG (Thermo Fisher Scientific A-11011).

#### Aortic pulse wave velocity (aPWV)

aPWV was assessed non-invasively using an Indus Doppler Flow Velocity System (Scintica, Canada) as previously described. Briefly, isoflurane (2%) was used to anesthetize mice that were placed supine with legs secured to ECG electrodes on a heated board. Doppler probes were placed on the skin at the transverse aortic arch and abdominal aorta, ∼ 4 cm apart. For each site, the pre-ejection time, or time between the R-wave of the ECG to the foot of the Doppler signal was determined. To calculate aPWV, the distance between the probes was divided by the difference in the thoracic and abdominal pre-ejection times and is presented as centimeters/second (cm/s). Following aPWV measures, mice were euthanized by exsanguination via cardiac puncture while anesthetized with isoflurane^30^.

#### Calcium content in aortas

Dried aorta was defatted with chloroform and methanol (2:1) for 48 hours and dehydrated with acetone for three hours. The dried samples were incinerated to ashes at 600°C for 24 hours using an electric muffle furnace (Thermo Scientific), then extracted with 0.6 N HCl. Calcium content from aortas was quantified using the o-cresolphthalein method. In addition, calcified lesions in the aortic arches were analyzed using von Kossa staining as previously described using a double-blind fashion as previously described^28-32^.

#### Statistical analysis

Data were collected from more than two independent experiments and reported as the means ± S.E.M. Statistical analysis for two-group comparison was performed using the Student’s *t* test, or one-way ANOVA or two-way ANOVA with a Newman-Keuls post-hoc test for multi-group comparison. Significance was accepted at *P* < 0.05.

### Study Approval

All animal protocols and experimental procedures were approved by the Institutional Committees at the University of Colorado-AMC.

## ACKNOWLWGEMENTS

This work is dedicated to the memory of Shinobu Miyazaki-Anzai, who sadly passed away before the completion of this project. This work was supported by grants from NIH HL132318, R01HL1157069 and R01DK124901 to M. Miyazaki. We would like to thank the personnel from the University of Colorado Anschutz Medical Campus Advanced Light Microscopy Core for training and use of the confocal and TIRF microscopes. Imaging was performed in the Advanced Light Microscopy Core Facility of the NeuroTechnology Center at the University of Colorado Anschutz Medical Campus, which is supported in part by a Rocky Mountain Neurological Disorders Core Grant (P30 NS048154) and by a Diabetes Research Center Grant (P30 DK116073).

---

**Supplemental Figure 1.**
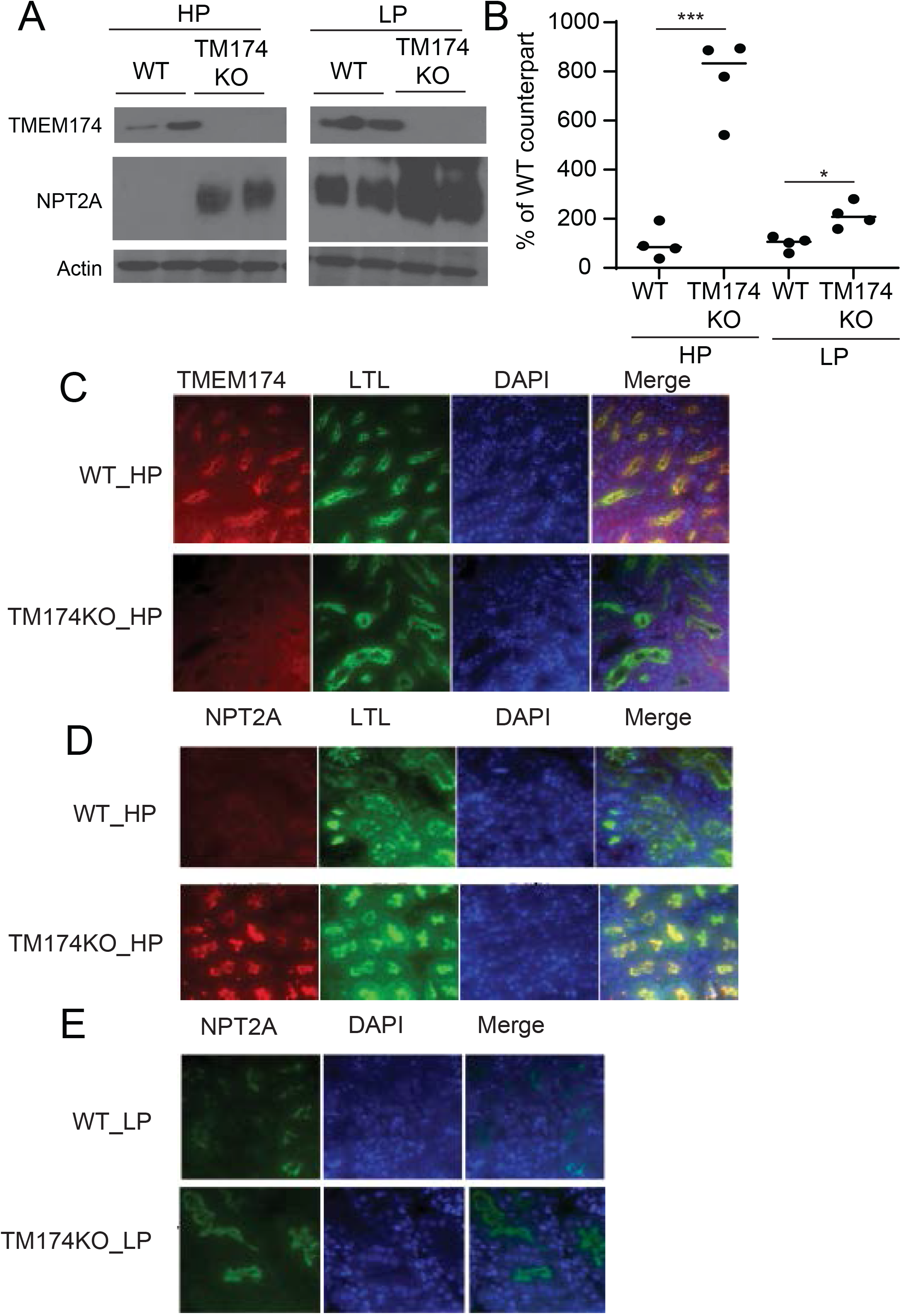
Low phosphate diet attenuates the effect of TMEM174 deficiency on NPT2A protein expression in the renal proximal tubules. A) Immunoblot and B) densitometry analyses of TMEM174 and NPT2A in the renal BBM from mice fed high phosphate and low phosphate diets. The renal BBM fraction was isolated using the Ca^2+^ precipitation method. Immunofluorescence analysis of C) TMEM174 and D) and E) NPT2A in TMEM174KO mice fed high-phosphate and low-phosphate diets. NPT2A and TMEM174 were detected with anti-mouse NPT2A and anti-mouse TMEM174 polyclonal antibodies coupled with LTL (a proximal tubule specific marker).

**Supplemental Figure 2.**
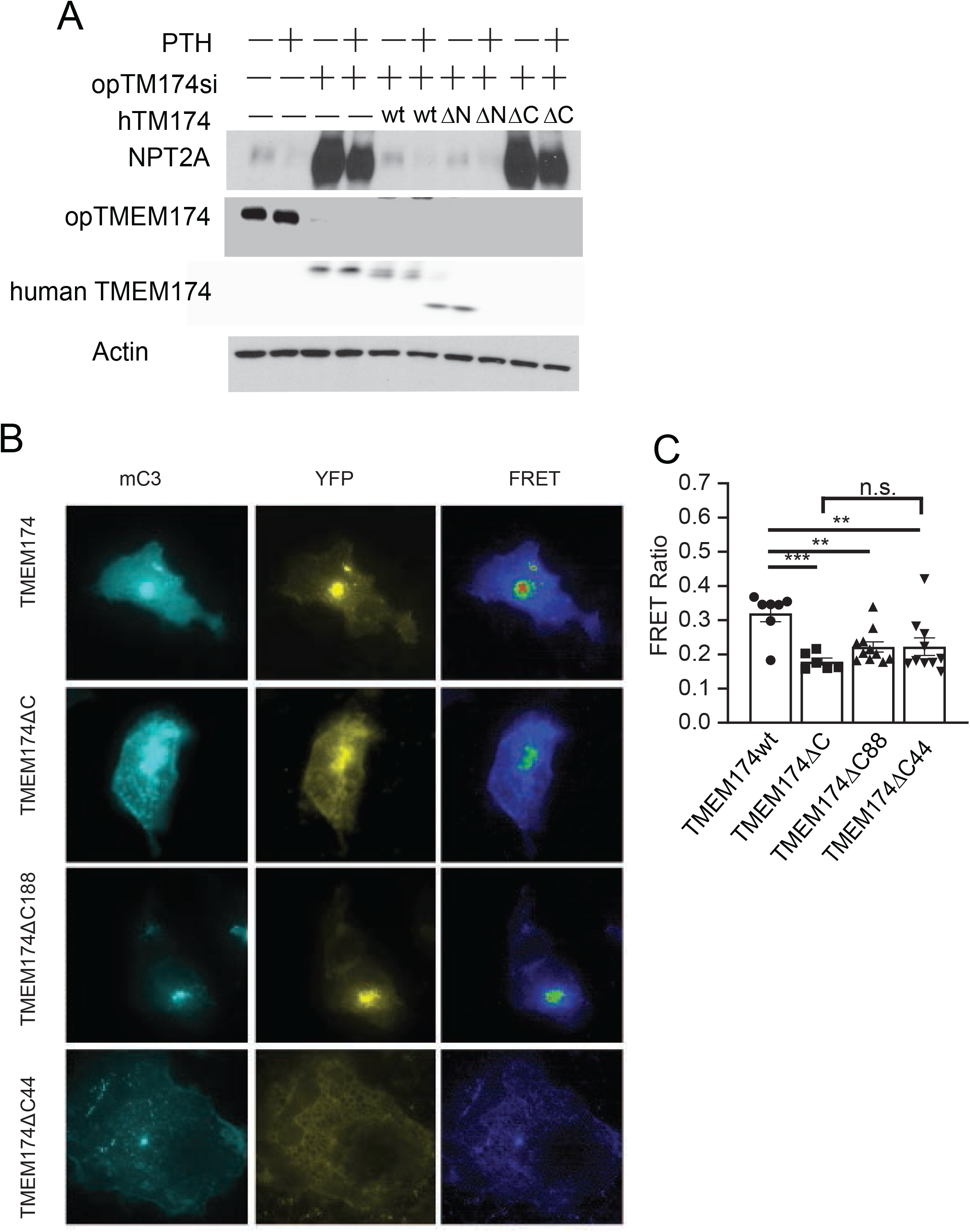
The TMEM174 C-terminal end is critical in the regulation of NPT2A. A) Immunoblot analysis of TMEM174 and NPT2A in TMEM174 knockdown OKP cells reconstituted with human TMEM174 mutants in the absence and presence of PTH. OKP cells were treated with opossum-specific TMEM174 for 7 days and then TMEM174 expression was reconstituted by human TMEM174 mutants. After two days at 100% confluency, cells were treated with 10nM PTH/0.1%BSA for 1 hour. Total cell lysate was isolated with RIPA buffer. Opossum and human TMEM174 were detected with anti-opossum TMEM174 specific antibody (Biomatik) and anti-FLAG monoclonal antibody (Sigma), respectively. B) The representative images for each set of experiments showing EYFP-NPT2A and TMEM174-mC3 wt and mutants lacking full length, 144 (ΔC), 86 (ΔC86) and 43 (ΔC43) amino acids of the TMEM174 C-terminal end. C) The bar graph represents the initial FRET ratio of experiments. When NPT2A is partnered with TMEM174wt it exhibits a higher FRET ratio compared to any TMEM174 in which C-terminal end truncations were made.

